# DOTA chelation through click chemistry enables favorable biodistribution of ^89^Zr-radiolabeled antibodies: A comparison with DFO chelation

**DOI:** 10.1101/2022.09.08.507067

**Authors:** Ryota Imura, Yoshitaka Kumakura, Lin Yan, Yuki Shimoura, Hiroyuki Takahashi, Hiroyuki Ida, Youichiro Wada, Nobuyoshi Akimitsu

**Author notes:** **Corresponding author** Yoshitaka Kumakura, M.D., Ph.D., Department of Diagnostic Radiology and Nuclear Medicine, Saitama Medical Center, Saitama Medical University, 1981 Kamoda, Kawagoe-shi, Saitama 350-8550 JAPAN.

## Abstract

Currently, the DFO chelator is commonly used to conjugate monoclonal antibodies (mAbs) and ^89^Zr, whereas the DOTA chelator is commonly used to conjugate mAbs and alpha- and beta-emitting metal radionuclides. However, if the degradation of [^89^Zr]Zr-DFO-mAb is not negligible, the in vivo biodistribution of ^89^Zr might not reflect that of metal radionuclides conjugated with DOTA-mAb. We hypothesized that [^89^Zr]Zr-DOTA-mAb as a new imaging counterpart would accurately predict the biodistribution of therapeutic metal radionuclides delivered by DOTA-mAb. In this study, we prepared [^89^Zr]Zr-DOTA-trastuzumab for the first time by a two-step reaction using click chemistry and then investigated the differences in biodistribution profiles between two chelating approaches for ^89^Zr.

**Methods:** We prepared [^89^Zr]Zr-DOTA-trastuzumab from DOTA-tetrazine conjugates (DOTA-Tz) and transcyclooctene-trastuzumab conjugates (TCO-trastuzumab). We first radiolabeled DOTA-Tz with ^89^Zr in a reaction solution of MeOH and HEPES buffer and then used a click reaction to obtain [^89^Zr]Zr-DOTA-Tz/TCO-trastuzumab. We performed biodistribution studies and PET imaging with [^89^Zr]Zr-DOTA-trastuzumab in a mouse model of HER2-positive ovarian cancer, SKOV3 xenograft mice at 24, 72, and 144 hours post-injection and compared these data with those of [^89^Zr]Zr-DFO-trastuzumab.

**Results:** TCO-trastuzumab was radiolabeled with [^89^Zr]Zr-DOTA-Tz in the two-step reaction in good radiochemical yield (57.8 ± 17.6%). HER2-positive tumors were clearly visualized with [^89^Zr]Zr-DOTA-trastuzumab in PET imaging studies. The temporal profile changes of ^89^Zr radioactivity in SKOV3 tumors and bone marrow were sufficiently different between [^89^Zr]Zr-DOTA-trastuzumab and [^89^Zr]Zr-DFO-trastuzumab (P < 0.05). Conclusion: [^89^Zr]Zr-DOTA-trastuzumab can be produced by the two-step radiolabeling reaction based on the Tz/TCO click reaction. Presumably, ^89^Zr released from DFO is not negligible. In contrast, [^89^Zr]Zr-DOTA-mAb would better predict the biodistribution of [^177^Lu]Lu- or [^225^Ac]Ac-DOTA-mAb than [^89^Zr]Zr-DFO-mAb, thus avoiding the use of different chelator for ^89^Zr at the expense of the click chemistry step.

**Graphical Abstract:** 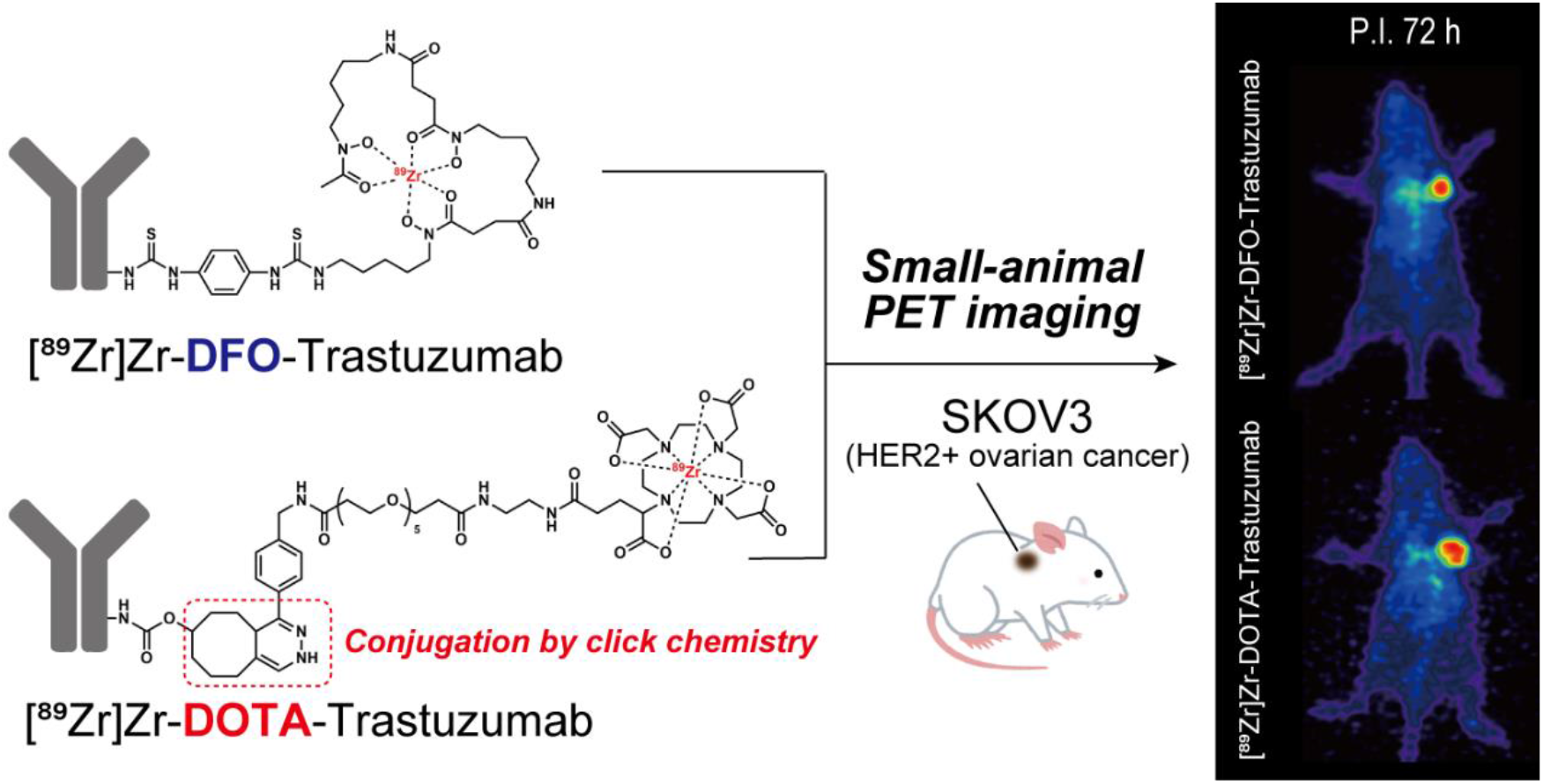

## Introduction

Monoclonal antibodies (mAbs) labeled with alpha and/or beta emitters are considered promising macromolecules for targeted radionuclide therapy (TRT) [1–3]. An imaging counterpart for TRT is also needed to visualize tumors [4,5]. Previous studies have used positron emitting ^64^Cu for tumor imaging [1–3]. However, the half-life of ^64^Cu (12.7 h) is not long enough to assess the in vivo distribution of long-circulating, slow-binding mAbs [1,6,7]. The half-life of positron emitting ^89^Zr (78.5 h) is suitable not only for visualization of tumors but also for assessment of cumulative radiation exposure to normal organs such as the bone marrow, liver, intestines and kidneys [1,5,7,8]. Currently, ^89^Zr radiolabeling for mAbs is often achieved via the DFO chelator, which reacts readily at room temperature [9–13], despite its known shortcomings (e.g. non-specific accumulation of free ^89^Zr in bones). As a direct substitute for [^177^Lu]Lu-DOTA-mAbs and [^225^Ac]Ac-DOTA-mAbs, ^89^Zr-labeled DOTA-mAbs would be preferable to replicate the therapeutic dose distribution. However, to the best of our knowledge, there is no report on the synthesis of [^89^Zr]Zr-DOTA-mAbs [14].

Recently, we developed a method for radiolabeling PSMA-617 containing DOTA with ^89^Zr at 90°C for 30 min in a mixture of HEPES buffer and organic solvents [15]. However, such direct radiolabeling cannot be used to prepare [^89^Zr]Zr-DOTA-mAbs due to irreversible thermal denaturation of mAbs.

To circumvent the denaturation of mAbs, we employed a two-step reaction with click chemistry [8]. Specifically, ^89^Zr was first coupled with DOTA at high temperature, followed by a click chemistry reaction in which [^89^Zr]Zr-DOTA was conjugated with trastuzumab at room temperature. We selected trastuzumab because it is one of the best studied mAbs for theranostic application. The aim of this study was to investigate in the delayed biodistribution of ^89^Zr between two chelating agents used for radiolabeling trastuzumab. There would be no measurable difference if the degradation products of [^89^Zr]Zr-DFO/DOTA-trastuzumab were similarly distributed in the organs. Therefore, we performed PET and ex vivo biodistribution studies in a mouse model of HER2-overexpressing human ovarian adenocarcinoma (mice with SKOV3 xenografts), and then determined the difference in biodistribution profiles between [^89^Zr]Zr-DOTA-trastuzumab and [^89^Zr]Zr-DFO-trastuzumab. Two-step reaction radiolabeling using click chemistry could be versatile and practical, and the use of [^89^Zr]Zr-DOTA would be suitable for numerous applications of mAbs.

## Materials and Methods

### Materials

The following chelating agents and click chemistry reagents were used: NH_2_-DOTA-GA (Chematech, France), tetrazine-PEG_5_-NHS ester (Click Chemistry Tools, USA), TCO-NHS ester (Click Chemistry Tools, USA), and p-SCN-Bn-deferoxamine (Macrocyclics, USA). SKOV3 cell line (ovarian cancer, ATCC HTB-77) was purchased from American Type Culture Collection (USA). Other reagents, buffers, or cell culture media were purchased from FUJIFILM Wako Pure Chemical (Japan), Dojindo Laboratories (Japan), or Sigma Aldrich (USA).

### Radiolabeling Experiments

We prepared [^89^Zr]Zr-DOTA-trastuzumab through a two-step reaction using click chemistry. Due to the high reaction rate (*k* ∼ 10^0^–10^6^ M^-1^ s^-1^) [16,17], we chose the inverse electron demand-Diels-Alder (IEDDA) cycloaddition between tetrazine (Tz) and transcyclooctene (TCO). The preparation and qualification of ^89^Zr followed a method described in the literature [18]. Briefly, ^89^Zr was produced by proton irradiation to ^89^Y targets, and the irradiated targets were purified to [^89^Zr]ZrCl_4_. The precursors (DOTA-Tz and TCO-trastuzumab) were synthesized as described in the supplementary material (Figure S1).

We first coupled DOTA-Tz with ^89^Zr and then conjugated TCO-trastuzumab with [^89^Zr]Zr-DOTA-Tz, as shown in Figure 1. In the first step, we added the following reagents to a 2-mL tube: 1 μL of DOTA-Tz solution (10^−2^ mol/L) in water, 499 μL of HEPES buffer (0.5 mol/L, pH 7.0), 1200 μL of methanol (MeOH), and 300 μL of purified ^89^Zr solution. The mixture was allowed to react at 90°C for 30 min. We evaluated the radiochemical yield by instant thin-layer chromatography (ITLC-SG, Agilent, USA) using 1 mol/L ammonium acetate and MeOH (1:1) as the mobile phase. We then removed the MeOH by nitrogen bubbling at 60°C for 10 min. In the second step, we allowed the TCO-trastuzumab (400 μg, 200 μL, 2000 μg/mL) to react in the [^89^Zr]Zr-DOTA-Tz solution at room temperature for 15 min. Then, the buffer of [^89^Zr]Zr-DOTA-trastuzumab was replaced with phosphate-buffered saline (PBS) using centrifugal filter units (Amicon Ultra 4, 50,000-Dalton molecular weight cutoff; Millipore, USA).

**Figure 1.**
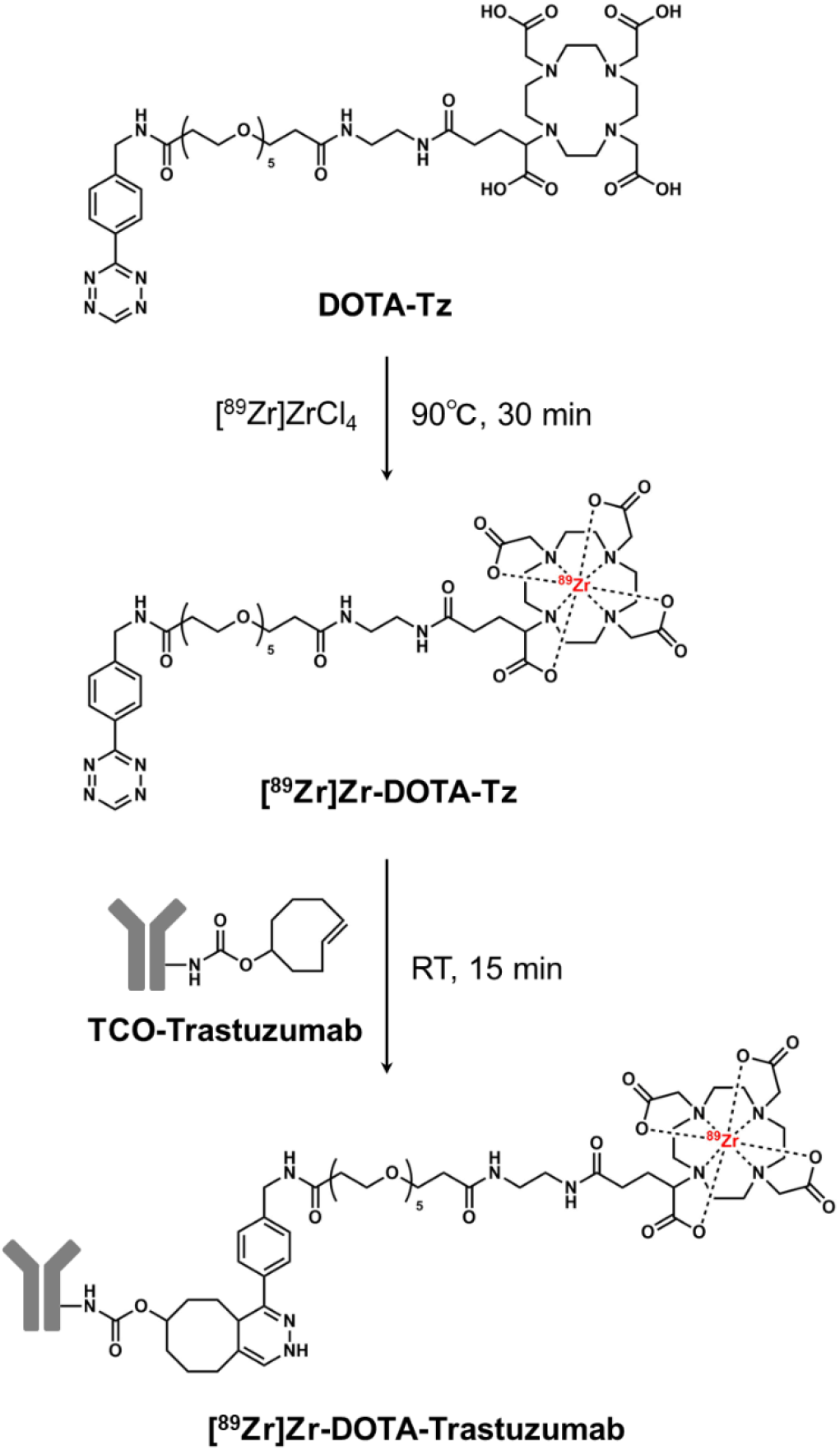
The novel two-step radiolabeling of [^89^Zr]Zr-DOTA-trastuzumab. ^89^Zr was first coupled with DOTA-Tz at 90°C for 30 min. TCO-trastuzumab was then conjugated at room temperature to give the final product [^89^Zr]Zr-DOTA-Tz/TCO-trastuzumab. It should be noted that trastuzumab was not heated at 90°C for 30 min.

We also prepared [^89^Zr]Zr-DFO-trastuzumab according to the literature [13,19] to perform comparisons with [^89^Zr]Zr-DOTA-trastuzumab. We prepared [^89^Zr]Zr-oxalate in 1 mol/L oxalic acid solution according to previous methods [11]. We added 100 μL of [^89^Zr]Zr-oxalate, 45 μL of 2 mol/L sodium carbonate solution, 655 μL of 0.5 mol/L HEPES buffer, and 200 μL of DFO-trastuzumab solution (2 mg/mL) to a microtube. The mixture was reacted at 25°C for 30 min (see the supplementary material for details).

### Cell Culture

We used the SKOV3 cell line, which highly expresses human epidermal growth factor receptor 2 (HER2), for in vitro assays and to create the mouse model for in vivo experiments. The SKOV3 cell line was cultured in Dulbecco’s modified Eagle medium (DMEM) supplemented with heat-inactivated 10% fetal calf serum. Cell culture was performed at 37°C in an atmosphere of 5% CO_2_. Cells were harvested with trypsin-ethylenediaminetetraacetic acid (trypsin-EDTA; 0.25% trypsin, 0.02% EDTA).

### Quality Control and In Vitro Assays

We performed the quality control of [^89^Zr]Zr-DOTA-trastuzumab and [^89^Zr]Zr-DFO-trastuzumab by size-exclusion chromatography (Figure S3). The in vitro stability of both radioligands was determined by EDTA challenge assays (Table S1). The immunoreactivity fractions of both radioligands were also examined by Lindmo assay (Figure S4) [20]. Experimental details of the size-exclusion chromatography, stability assays, and Lindmo assays can be found in the Supplementary Materials.

### Mouse Tumor Model

This study was approved by the Animal Experimentation Committee of the Isotope Science Center, the University of Tokyo. We performed all animal experiments in accordance with the University Animal Experimentation Regulations and the guidelines of ARRIVE.

Seven-week-old female nude mice (BALB/c *nu/nu*) were purchased from Japan SLC Inc. The mice received a subcutaneous injection of SKOV3 cells (5 × 10^6^ cells, 100 μL) suspended in 50% Matrigel (Corning, USA) into the right shoulder. We used this mouse tumor model for biodistribution and PET imaging studies.

### Biodistribution Studies

To examine the ex vivo biodistribution of [^89^Zr]Zr-DFO-trastuzumab and [^89^Zr]Zr-DOTA-trastuzumab, mice were injected with each radioligand (∼5 μg, ∼0.1 MBq per mouse) via the tail vein and were sacrificed at 24, 72, or 144 h after injection (N = 4 at each time point and for each ligand). Organs of interest (blood, liver, spleen, kidney, stomach, large intestine, small intestine, heart, lung, tumor, muscle, bone, and skin) were dissected and weighed. The radioactivity of each organ was immediately measured with a gamma counter (Cobra Quantum, Perkin Elmer) and calculated as %ID/g.

### Statistical Analysis

All data were expressed as mean and standard deviation. To analyze the biodistribution profiles of each radioligand, we compared the data of [^89^Zr]Zr-DFO-trastuzumab and [^89^Zr]Zr-DOTA-trastuzumab for each organ at each time point (24, 72, and 144 h after injection) using two-way analysis of variance (ANOVA) of GraphPad Prism 7. In the two-way ANOVA, we examined the effects of chelators, time, and their interaction on the radioactivity accumulation in the organs (%ID/g). P values less than 0.05 were considered statistically significant.

### PET Imaging Studies

To identify unfavorable accumulation in the organs, PET imaging studies with [^89^Zr]Zr-DFO-trastuzumab and [^89^Zr]Zr-DOTA-trastuzumab were performed 24, 72, and 144 h after the dose injection (∼50 μg, ∼ 3 MBq per mouse) via the tail vein. We used a Clairvivo PET scanner (Shimadzu, Japan) and SKOV3 tumor mice (N = 4 for each chelator group), which were not used in the biodistribution studies. The mice were under isoflurane anesthesia during the 30 min PET scans.

## Results

### Radiolabeling Experiments

We successfully prepared [^89^Zr]Zr-DOTA-trastuzumab for the first time by a two-step reaction. Mass spectrometry confirmed the successful synthesis of the precursors TCO-trastuzumab (Figure S2) and DOTA-Tz (Supplementary Material 1.2). The radiochemical yield (RCY) of [^89^Zr]Zr-DOTA-Tz was 59.3 ± 14.9% and that of [^89^Zr]Zr-DOTA-trastuzumab was 57.8 ± 17.6%. The results of size-exclusion chromatography (Figure S3) showed that the high radiochemical purity of [^89^Zr]Zr-DOTA-trastuzumab (> 95%) was achieved. The results of the EDTA challenge assays (Table S1) showed that [^89^Zr]Zr-DOTA-trastuzumab was stable even in excess amount of EDTA (> 90%). The percentage of immunoreactivity determined by the Lindmo assay was 95% (Figure S4(b)).

Likewise, high radiochemical purity over 95% (Figure S3), high stability in excess EDTA (> 90%, Table S1), and high immunoreactivity (98%, Figure S4(b)) were confirmed in the preparation of [^89^Zr]Zr-DFO-trastuzumab.

### Biodistribution Studies

[^89^Zr]Zr-DFO-trastuzumab and [^89^Zr]Zr-DOTA-trastuzumab were then used for biodistribution studies SKOV3 tumor-bearing mice, with timepoints of 24, 72, and 144 h after injection (N = 4 at each time point). The accumulation of [^89^Zr]Zr-DFO-trastuzumab in the tumor increased over time, and the highest accumulation (31.1 ± 12.3%ID/g) was observed 144 h after injection, which was 2.1 times higher than that at 72 h (14.7 ± 2.3% ID/g). In contrast, the accumulation of [^89^Zr]Zr-DOTA-trastuzumab in the tumor peaked 72 h after injection (30.8 ± 7.3%ID/g) and then decreased at 144 h after injection (24.5 ± 8.3%ID/g). Notably, the accumulation of [^89^Zr]Zr-DOTA-trastuzumab in the bone decreased over time, whereas that of [^89^Zr]Zr-DFO-trastuzumab increased.

Using the two-way ANOVA, we found that the radioactivity uptake in kidney, bone, and skin differed significantly depending on the chelators (DFO versus DOTA); [^89^Zr]Zr-DOTA-trastuzumab accumulated less in kidney, bone, and skin than [^89^Zr]Zr-DFO-trastuzumab at each time point (24, 72, and 144 h after injection). We combined the ex vivo biodistribution data of Figures 2a and 2b, and then reorganized them to show the time-%ID/g curves of two chelators for each organ, as shown in Figure S5. We also found a significant interaction in tumor (P = 0.0239) and bone (P = 0.0104) uptake between the radioligands ([^89^Zr]Zr-DFO-trastuzumab and [^89^Zr]Zr-DOTA-trastuzumab). [^89^Zr]Zr-DFO-trastuzumab and [^89^Zr]Zr-DOTA-trastuzumab showed different temporal changes in the tumor and bone. We found no significant interactions in the uptake between the two radioligands at the three time points assessed in the other tissues (P > 0.05). All P values obtained in the two-way ANOVA are shown in Table S3.

**Figure 2.**
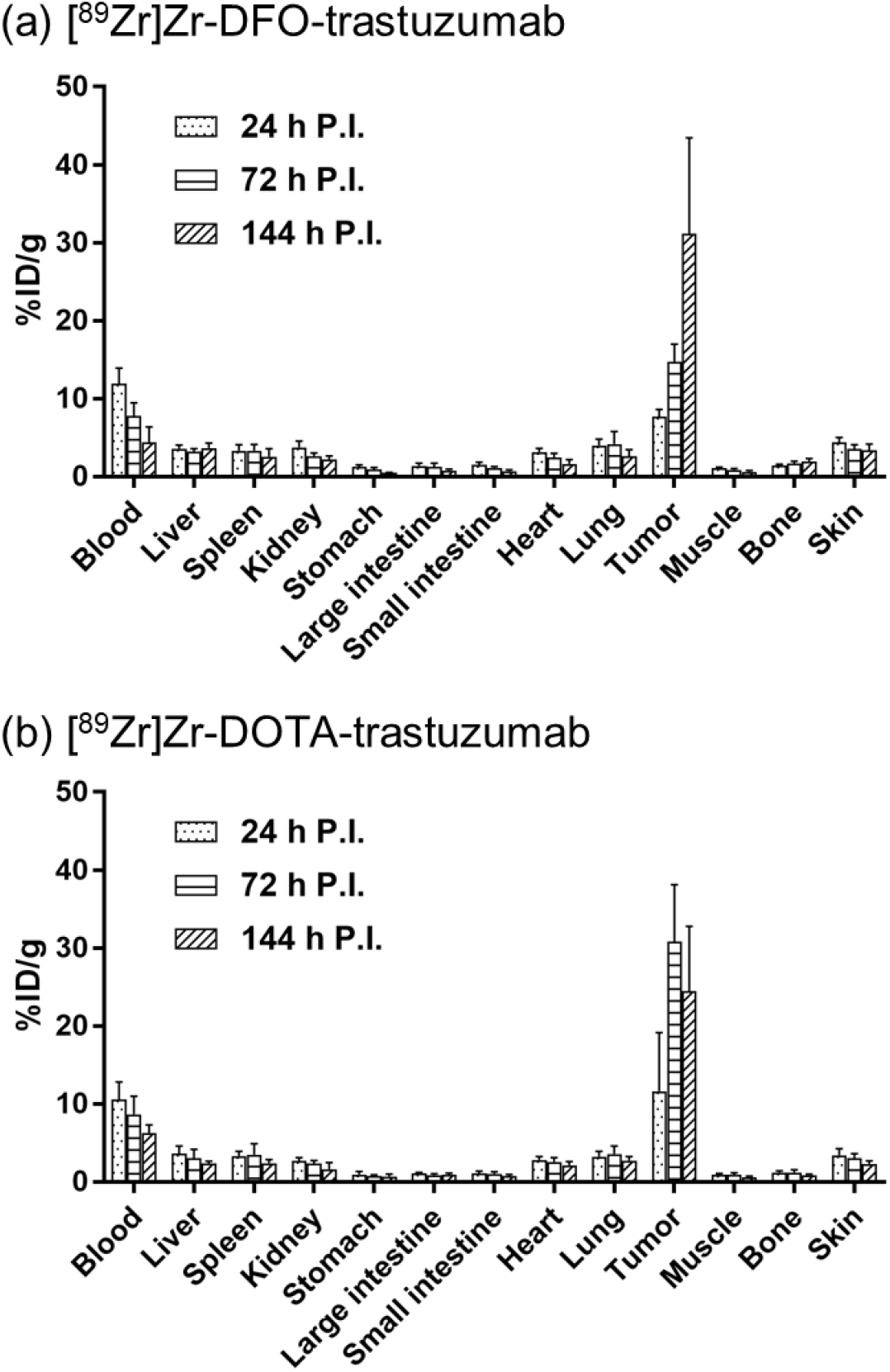
Biodistribution profiles in SKOV3 tumor-bearing mice at 24, 72, and 144 h postinjection for (a) [^89^Zr]Zr-DFO-trastuzumab (N = 4 at each time point) and (b) [^89^Zr]Zr-DOTA-trastuzumab (N = 4 at each time point). Error bars indicate standard deviations. Each dose per mouse was ∼ 5 μg (∼0.1 MBq). Organs of interest were dissected and weighed to calculate %ID/g.

### PET Imaging Studies

Using mice bearing HER2 positive tumors, we compared PET images of [^89^Zr]Zr-DOTA-trastuzumab with that of [^89^Zr]Zr-DFO-trastuzumab. Figure 3 shows representative maximum intensity projection (MIP) PET images up to six days (144 h) postinjection. No noticeable differences in tumor accumulation was observed. No unexpected accumulation was observed in the organs. [^89^Zr]Zr-DOTA-trastuzumab was useful for PET imaging to clearly identify tumor size and location.

**Figure 3.**
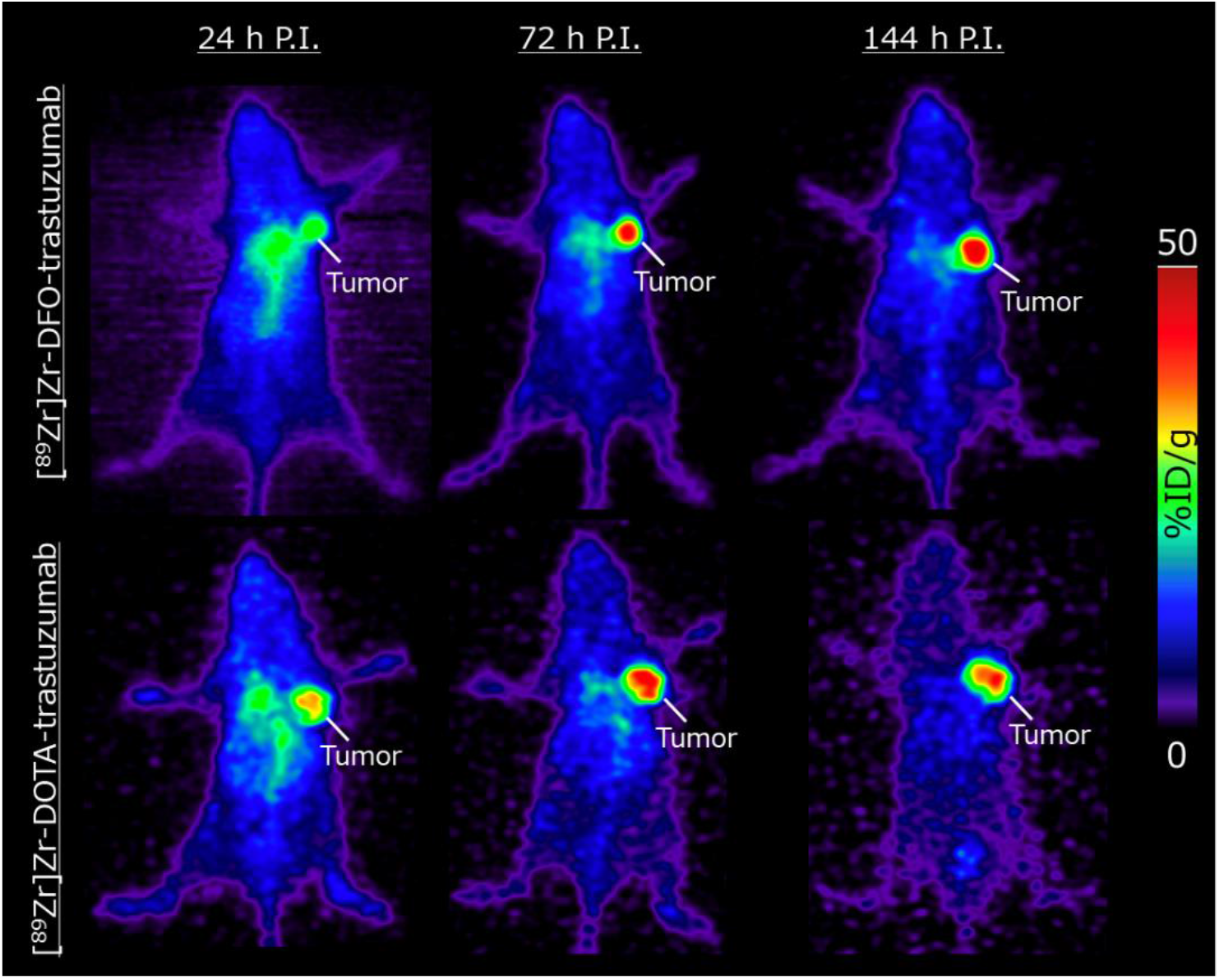
Representative maximum intensity projection (MIP) PET images of [^89^Zr]Zr-DFO-trastuzumab and [^89^Zr]Zr-DOTA-trastuzumab using SKOV3 tumor-bearing mice at 24, 72, and 144 h postinjection. Each dose per mouse was ∼50 μg (∼3 MBq), and the mice were under isoflurane anesthesia for 30 min PET scanning.

## Discussion

Monoclonal antibodies labeled with radionuclides through DOTA provide numerous theranostic options. To radiolabel trastuzumab for small animal PET imaging of HER2-expressing xenografts, we used the novel combination of ^89^Zr and DOTA and compared it with the combination of ^89^Zr and DFO. The long half-life (3.2 d) of ^89^Zr is attractive for PET imaging of trastuzumab, creating the ideal companion diagnostic agent to be used prior to TRT with ^177^Lu and ^225^Ac. However, the incorporation of ^89^Zr with DOTA requires heating, which can denature trastuzumab. In this study, we circumvented this problem by introducing a two-step reaction using click chemistry (inverse electron demand-Diels-Alder reaction), established a radiolabeling method, and obtained a high yield of [^89^Zr]Zr-DOTA-trastuzumab (RCY of [^89^Zr]Zr-DOTA-Tz: 59.3 ± 14.9%, RCY of [^89^Zr]Zr-DOTA-trastuzumab: 57.8 ± 17.6%). Although tetrazine is known to degrade in FBS at 37°C [21], we have demonstrated that [^89^Zr]Zr-DOTA-Tz reacted well with TCO-trastuzumab after the radiolabeling reaction of 90°C for 30 min in HEPES buffer and MeOH. By establishing the method to prepare [^89^Zr]Zr-DOTA-Tz, we have synthesized [^89^Zr]Zr-DOTA-trastuzumab for the first time.

Our small animal PET experiments successfully visualized HER2-expressing SKOV3 tumors. Having compared DOTA with DFO, we showed the different temporal changes in ^89^Zr radioactivity of the tumor and bone between the two chelators. The tumor accumulation of [^89^Zr]Zr-DFO-trastuzumab continued to increase until the end of the biodistribution experiments (Figure 2 and Figure S5(l)), while that of DOTA-conjugated trastuzumab increased to a peak at 72 h postinjection (Figure 2 and Figure S5(j)). This peak at 72 h agrees with the previously reported peak of [^177^Lu]Lu-DOTA-trastuzumab [22]. This notable difference could be explained by two putative factors. First, the degradation of [^89^Zr]Zr-DFO-trastuzumab released free ^89^Zr, which ultimately accumulated in the bone marrow [23,24]. Second, the degradation of [^89^Zr]Zr-DOTA-trastuzumab generated [^89^Zr]Zr-DOTA, which did not release free ^89^Zr. This phenomenon was probably due to the strong DOTA incorporation of ^89^Zr ions as well as the rapid renal clearance of plasma [^89^Zr]Zr-DOTA into urine [25]. Thus, the postdegradation forms of ^89^Zr are also important in understanding the difference in ^89^Zr biodistribution and its temporal changes. Larger ^89^Zr uptake in kidney and skin for [^89^Zr]Zr-DOTA-trastuzumab compared with that for [^89^Zr]Zr-DFO-trastuzumab might be due to non-specific accumulation of free ^89^Zr.

The difference in the temporal profile changes of HER2-expressing SKOV3 tumors could also be explained by the differences in the degradation processes between [^89^Zr]Zr-DFO-trastuzumab and [^89^Zr]Zr-DOTA-trastuzumab. Due to the weak metal ion incorporation of DFO, free ^89^Zr released in plasma might have been captured by intratumoral proteins in a manner similar to the accumulation mechanism of [^67^Ga]Ga citrate. [26,27]. For this reason, the tumor accumulation of [^89^Zr]Zr-DFO-trastuzumab might continue to increase, while that of [^89^Zr]Zr-DOTA-trastuzumab continued to decrease.

The limitation of this study is that our [^89^Zr]Zr-DOTA-trastuzumab contains the Tz/TCO structure, which is presumably not degradable in vivo, while [^177^Lu]Lu-DOTA-mAb generally does not contain this structure [22]. The effects of the Tz/TCO structure on biodistribution must be investigated in future research using metal radionuclides other than ^89^Zr. Alternatively, this two-step radiolabeling method could be extended to label mAbs with alpha and beta emitters since a preparation method of M-DOTA-Tz (M = ^90^Y, ^177^Lu, ^225^Ac) has already been established [28–30].

## Conclusion

This study established a preparation method for [^89^Zr]Zr-DOTA-trastuzumab and performed PET imaging studies for the first time. We showed that [^89^Zr]Zr-DOTA-trastuzumab can be used to visualize HER2-positive tumors in small animals and may be a better imaging agent for [^177^Lu]Lu- or [^225^Ac]Ac-DOTA-mAb than [^89^Zr]Zr-DFO-trastuzumab because of the use of a common chelator.

## Abbreviations

DOTA: 1,4,7,10-tetraazacyclododecane-1,4,7,10-tetraacetic acid
DFO: deferoxamine B

## Acknowledgments

We gratefully thank Toshifumi Omura (JFE Engineering Corporation) and Toru Matsumoto (JFE Engineering Corporation) for technical help.

## Competing interests

Ryota Imura and Hiroyuki Ida are employees of JFE Engineering Corporation. This research was partially carried out with research funds from JFE Engineering Corporation. No other potential conflicts of interest relevant to this article exist.

## Notes

Financial support: This research was carried out with research funds from JFE Engineering Corporation, which employs the authors Ryota Imura and Hiroyuki Ida.

### Competing Interest Statement

Ryota Imura and Hiroyuki Ida are employees of JFE Engineering Corporation. This research was partially conducted with research funds of the JFE Engineering Corporation.

