## supplemental material for "DOTA chelation through click chemistry enables favorable biodistribution of ^89^Zr-radiolabeled antibodies: A comparison with DFO chelation"

### Supplementary Material

#### 1. Synthesis of DOTA-Tz

##### 1.1 Materials and Methods

DOTA-Tz was synthesized by amidation from $NH_2$-DOTA-GA and a tetrazine-$PEG_5$-NHS ester, as shown in Figure S1. $NH_2$-DOTA-GA (38.0 mg, 0.073 mmol), tetrazine-$PEG_5$-NHS ester (44.3 mg, 0.073 mmol), and triethylamine (0.021 mL, 0.15 mmol) were added to 0.76 mL of dimethylformamide (DMF) and then reacted at room temperature overnight. DOTA-Tz was fractionated by preparative high-performance liquid chromatography (HPLC) with a YMC-Actus Triart C18 column (φ 4.6 mm × 50 mm, 5.0 μm) using water containing 0.1% tetrafluoroacetic acid (solvent A) and acetonitrile (solvent B) in a 5–95% solvent B gradient over 30 min at a flow rate of 2 mL/min. The fractionated DOTA-Tz was then lyophilized, and the molecular weight of the synthesized compound was measured by LC-MS.

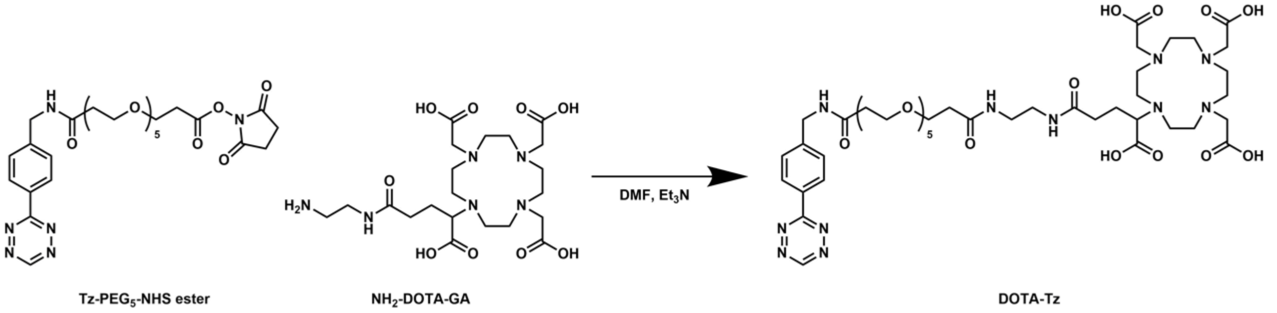


Figure S1. Synthesis of DOTA-Tz from Tz-$PEG_5$-NHS ester and $NH_2$-DOTA-GA.

#### 1.2 Results

DOTA-Tz was successfully obtained as a vivid pink powder. The yield was 77%. The *m*/*z* determined by mass spectrometry was 1008.25, which is consistent with the L + $H^+$ ion ($C_{44}H_{70}N_{11}O_{16}^+$, *m*/*z* = 1008.50). The obtained DOTA-Tz powder was stored at −80 °C.

#### 2. Preparation of DFO-Trastuzumab and TCO-Trastuzumab

#### 2.1 Materials and Methods

DFO-conjugated trastuzumab (DFO-trastuzumab) was prepared by using amine-reactive p-SCN-Bn-deferoxamine. Trastuzumab (2 mg) was dissolved in 0.1 mol/L $NaHCO_3$ buffer (pH 9.0), and the stock solution of p-SCN-Bn-deferoxamine in DMSO (10 mg/mL) was added to the mAb solution to yield a DFO:mAb reaction stoichiometry of 10:1. The resulting solution was incubated with gentle shaking for 1 h at 37 °C. Next, the buffer containing the modified antibody was replaced with PBS by using centrifugal filter units (Amicon Ultra 4, 50 kDa MWCO, Merck Millipore). The prepared DFO-trastuzumab was stored at -80 °C.

TCO-conjugated trastuzumab (TCO-trastuzumab) was prepared by the same method as that used for DFO-mAb except that TCO-NHS ester (Click Chemistry Tools) was used instead of p-SCN-Bn-deferoxamine.

The chelator-to-antibody ratio (CAR) of DFO-trastuzumab and TCO-trastuzumab was estimated by LC–MS measurements. Before the LC–MS measurements, glycans, which have different molecular weights for each antibody, were removed by PNGaseF (P705S, New England Biolabs) to prevent mass spectral splitting. Mass spectrometry was conducted with an Orbitrap Q Exactive (Thermo Fisher) instrument, and the data were deconvoluted to zero-charge mass. Unmodified trastuzumab were performed as a reference.

##### 2.2 Results

Figure S2 shows the deconvoluted zero-charge mass distribution of DFO-trastuzumab and TCO-trastuzumab prepared in this study. There are several mass peaks corresponding to the DFO- or TCO-conjugated mAbs because the mass intervals between each peak almost corresponded to the molecular weight of the payloads. The average CAR of DFO-trastuzumab was estimated as 1.11 from the relative intensity of each mass peak, and the average CAR of TCO-trastuzumab was estimated as 1.52.

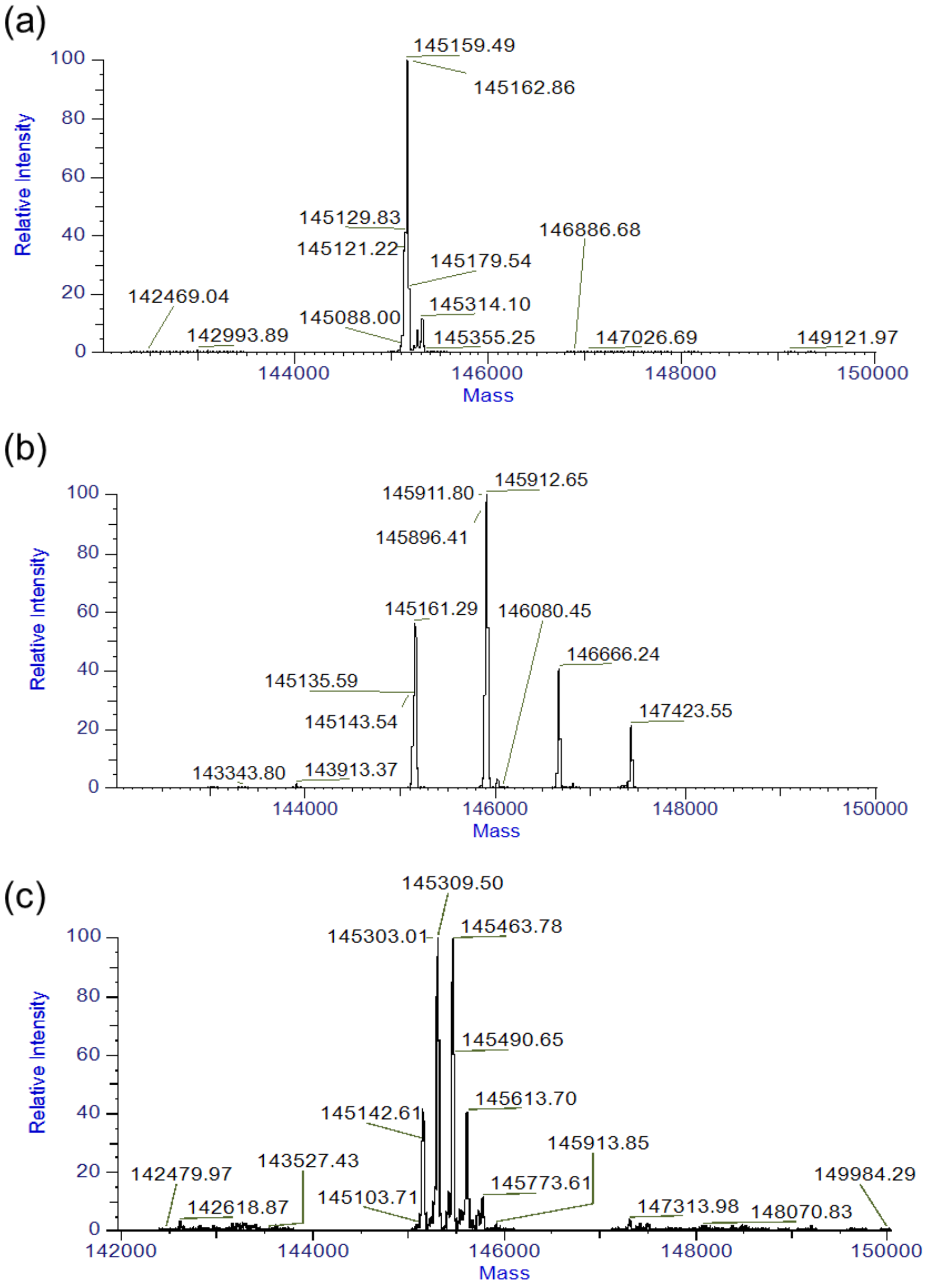


Figure S2. Deconvoluted zero-charge mass distributions obtained in LC-MS measurements. (a) Mass distribution of trastuzumab. Only a mass peak corresponding to CAR = 0 was observed (~145200 Da). (b) Mass distribution of DFO-trastuzumab. There are four mass peaks that correspond to CARs of 0 to 3. The intervals between each peak almost corresponded to the molecular weight of p-SCN-Bn-deferoxamine (752 g/mol). (c) Mass distribution of TCO-trastuzumab. There are four mass peaks that correspond to CARs of 0 to 4. The intervals between each peak almost corresponded to the molecular weight of TCO (152 g/mol).

#### 3. HPLC Analysis of $^{89}$Zr-Labeled Trastuzumab

##### 3.1 Materials and Methods

Quality control of [$^{89}$Zr]Zr-DFO-trastuzumab and [$^{89}$Zr]Zr-DOTA-trastuzumab was performed by size-exclusion chromatography (COSMOSIL 5Diol-300-II Packed Column, 7.5 mml.D.×300 mm, NACALAI TESQUE). The measurements were performed using radio-HPLC with 100 mmol/L sodium phosphate (pH 6.2–7.0) buffer including 150 mmol/L sodium chloride and 10 mmol/L sodium azide at a flow rate of 1.0 mL/min. HPLC was performed with a 280 nm UV absorbance detector and a radioactivity detector.

##### 3.2 Results

Figure S3 shows the size-exclusion chromatography results of [$^{89}$Zr]Zr-DFO-trastuzumab and [$^{89}$Zr]Zr-DOTA-trastuzumab. The aggregation peaks of each ligand were small.

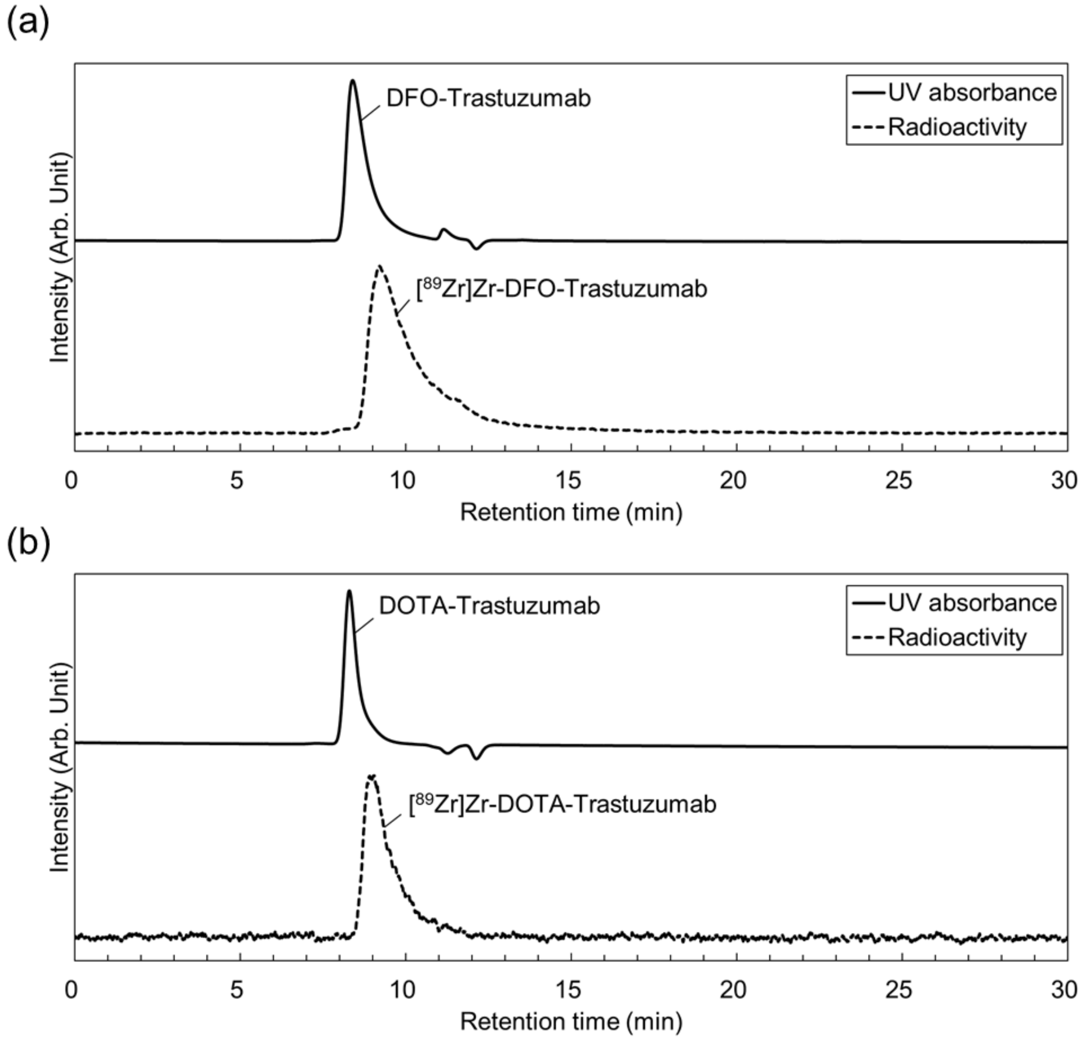


Figure S3. Size-exclusion chromatography analysis results of (a) [$^{89}$Zr]Zr-DFO-trastuzumab and (b) [$^{89}$Zr]Zr-DOTA-trastuzumab. The measurements were performed using radio-HPLC equipped with a 280 nm UV absorbance detector and a radioactivity detector and both data was collected at same time.

#### 4. <u>Evaluation of in vitro Stability</u>

##### 4.1 Materials and Methods

The in vitro stability of [$^{89}$Zr]Zr-DFO-trastuzumab and [$^{89}$Zr]Zr-DOTA-trastuzumab was first assessed by EDTA challenge tests. Labeled trastuzumab was added to microtubes and diluted to 200 μg/mL with PBS. Then, 1,000 mol-fold EDTA was added. The mixture was incubated at 37 °C. Aliquots were analyzed by ITLC-SA after 24, 72 and 144 h. This experiment was conducted in triplicate.

Next, the stability of the labeled trastuzumab in murine serum (FUJIFILM Wako Chemical, Japan) was also evaluated. Radiolabeled trastuzumab (200 μg/mL, 100 μL) and 500 μL of murine serum were added to microtubes. Then, the mixture was incubated at 37 °C. ITLC measurements were also performed at the same time points as the EDTA challenge tests. All assays were conducted triplicate.

**4.2 Results**

TABLE S1 shows the stability of [$^{89}$Zr]Zr-DFO-trastuzumab and [$^{89}$Zr]Zr-DOTA-trastuzumab in the EDTA challenge study at 37 °C and pH 7 over 7 days. TABLE S2 shows the stability of [$^{89}$Zr]Zr-DFO-trastuzumab and [$^{89}$Zr]Zr-DOTA-trastuzumab in murine serum. According to the results of the in vitro stability tests, [$^{89}$Zr]Zr-DFO-trastuzumab and [$^{89}$Zr]Zr-DOTA-trastuzumab showed high stability for 7 days (over 90%).

TABLE S1. Stability of [$^{89}$Zr]Zr-DFO-trastuzumab and [$^{89}$Zr]Zr-DOTA-trastuzumab in EDTA challenge tests at 37 °C and pH 7 over 7 days.

| Radioligand | % Intact | | |
|---|---|---|---|
| | 24 h | 72 d | 144 d |
| [$^{89}$Zr]Zr-DFO-mAb | 92.5 ± 0.6 | 97.9 ± 0.2 | 97.8 ± 0.3 |
| [$^{89}$Zr]Zr-DOTA-mAb | 94.0 ± 0.5 | 95.4 ± 0.6 | 95.0 ± 0.5 |

TABLE S2. Stability of [$^{89}$Zr]Zr-DFO-trastuzumab and [$^{89}$Zr]Zr-DOTA-trastuzumab in murine serum at 37 °C and pH 7 for 7 days.

| Radioligand | % Intact | | |
|---|---|---|---|
| | 24 h | 72 d | 144 d |
| [$^{89}$Zr]Zr-DFO-mAb | 98.2 ± 0.2 | 97.3 ± 0.5 | 95.9 ± 0.6 |
| [$^{89}$Zr]Zr-DOTA-mAb | 93.3 ± 2.0 | 94.6 ± 1.9 | 94.8 ± 1.2 |

#### 5. Lindmo Assay

##### 5.1 Materials and Methods

The immunoreactivity of [$^{89}$Zr]Zr-DFO-trastuzumab and [$^{89}$Zr]Zr-DOTA-trastuzumab was evaluated by the Lindmo assay [3]. An aliquot of radiolabeled trastuzumab ($10^{-10}$ mol/L) was added to SKOV3 cells suspended in DMEM containing 1% bovine serum albumin (BSA) ($1.6 \times 10^5 – 2.0 \times 10^7$

cells/mL). The cells were incubated for 1 h at 4 °C with continuous mixing throughout the incubation period to keep the cells in suspension. The cells and supernatant were separated by filtration. The cells on the filters were washed twice with PBS. The washed cells were measured in a gamma counter (Hidex), and the fraction of SKOV3 cells with bound radiolabeled trastuzumab was calculated (cell-binding fraction). The inverse of the cell-binding fraction was plotted against the inverse of the SKOV3 cell concentration. The y-intercept of the plot was calculated by linear regression analysis using GraphPad Prism 7.

**5.2 Results**

The results of the Lindmo assay are shown in Figure S4. The immunoreactivity of [$^{89}$Zr]Zr-DFO-trastuzumab was 98% and that of [$^{89}$Zr]Zr-DOTA-trastuzumab was 95%. Both radioligands prepared in this study were found to possess high immunoreactivity.

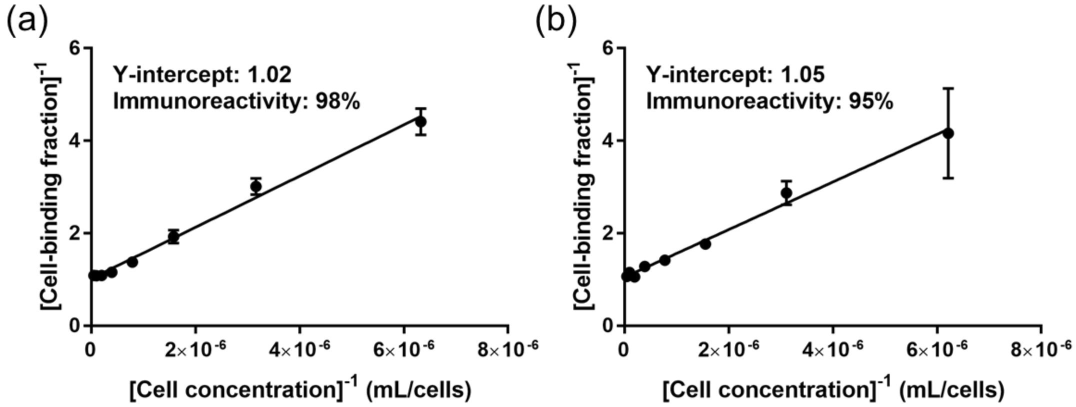


Figure S4. The plots of inverse of cell-binding fraction against inverse of SKOV3 cell concentration to assess the immunoreactivity of each radioligand. (a) For [$^{89}$Zr]Zr-DFO-trastuzumab, the y-intercept of the plot was 1.02 ± 0.05, and immunoreactivity was determined to be above 93%. (b) For [$^{89}$Zr]Zr-DOTA-trastuzumab, the y-intercept of the plot was 1.05 ± 0.05, and immunoreactivity was determined to be above 91%.

#### 6. Temporal Biodistribution Profiles

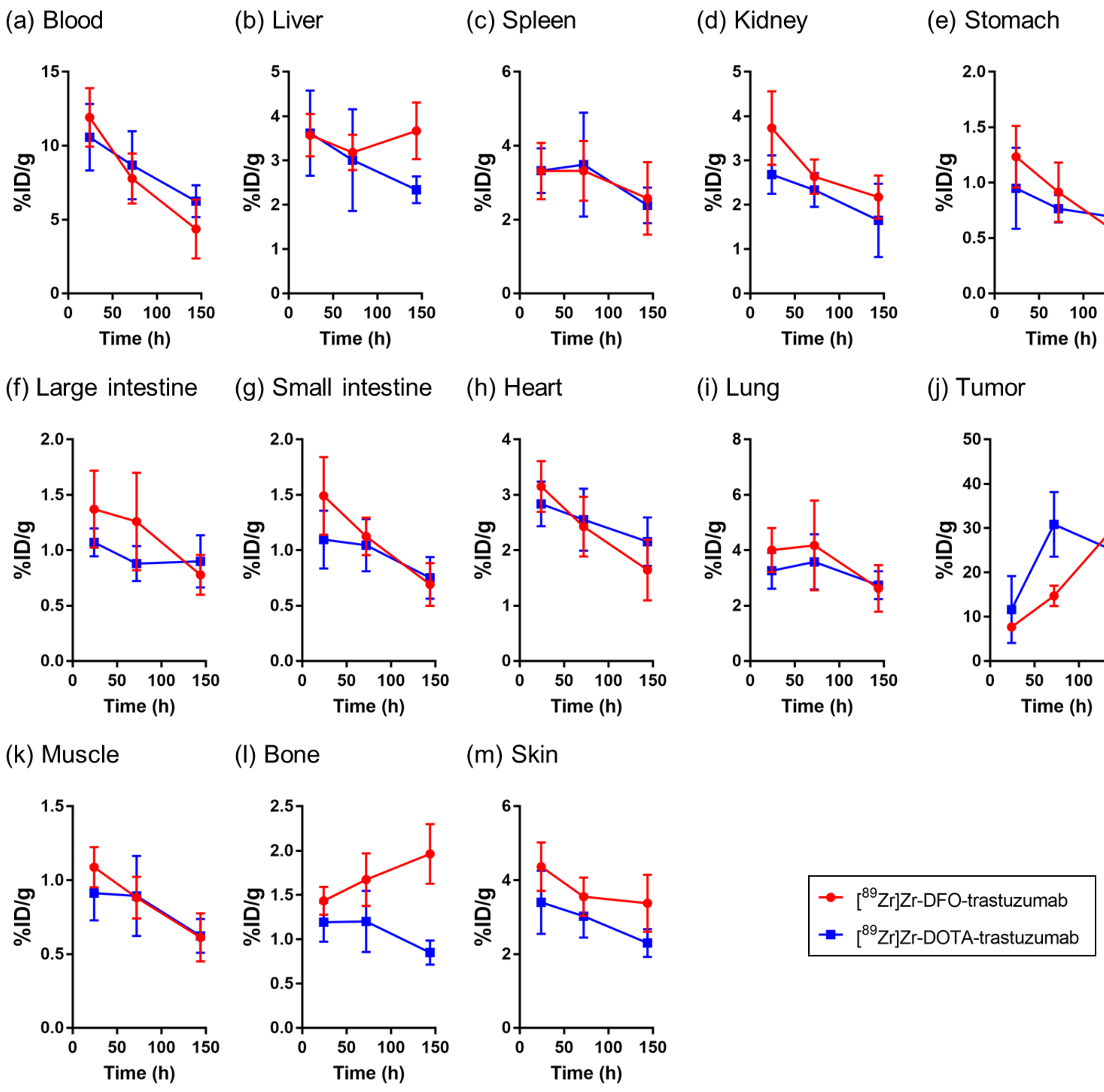


Figure S5. The time-%ID/g curves of two chelators for each organ. The value of %ID/g of Figure 2a and 2b in the main text are combined for each organ to make it easier to see the temporal changes between the two radioligand.

TABLE S3. P values obtained by the two-way ANOVA in the temporal profiles of the biodistribution ([$^{89}$Zr]Zr-DFO-trastuzumab versus [$^{89}$Zr]Zr-DOTA-trastuzumab). P values more than 0.05 are summarized by "ns", less than 0.05 by *, less than 0.01 by **, less than 0.001 by ***, and less than 0.0001 by ****.

| P value (summary) | Source of variance | | |
|---|---|---|---|
| | Effects of chelator | Time | Interaction |
| Blood | 0.5519 (ns) | <0.0001 (****) | 0.2563 (ns) |
| Liver | 0.1170 (ns) | 0.2417 (ns) | 0.1522 (ns) |
| Spleen | 0.9898 (ns) | 0.1014 (ns) | 0.9265 (ns) |
| Kidney | 0.0185 (*) | 0.0014 (**) | 0.4505 (ns) |
| Stomach | 0.4045 (ns) | 0.0040 (**) | 0.2227 (ns) |
| Large intestine | 0.1118 (ns) | 0.0360 (*) | 0.1689 (ns) |
| Small intestine | 0.1762 (ns) | 0.0006 (***) | 0.1849 (ns) |
| Heart | 0.6066 (ns) | 0.0013 (**) | 0.2724 (ns) |
| Lung | 0.3181 (ns) | 0.0559 (ns) | 0.6433 (ns) |
| Tumor | 0.1607 (ns) | 0.0004 (***) | 0.0239 (*) |
| Muscle | 0.4824 (ns) | 0.0012 (**) | 0.4908 (ns) |
| Bone | <0.0001 (****) | 0.6244 (ns) | 0.0104 (*) |
| Skin | 0.0045 (**) | 0.0159 (*) | 0.6781 (ns) |
